# Collection of 2429 constrained headshots of 277 volunteers for deep learning

**DOI:** 10.1101/2020.10.14.337220

**Authors:** Saki Aoto, Mayumi Hangai, Hitomi Ueno-Yokohata, Aki Ueda, Maki Igarashi, Yoshikazu Ito, Motoko Tsukamoto, Tomoko Jinno, Mika Sakamoto, Yuka Okazaki, Fuyuki Hasegawa, Hiroko Ogata-Kawata, Saki Namura, Kazuaki Kojima, Masao Kikuya, Keiko Matsubara, Kosuke Taniguchi, Kohji Okamura

## Abstract

Deep learning has rapidly been filtrating many aspects of human lives. In particular, image recognition by convolutional neural networks has inspired numerous studies in this area. Hardware and software technologies as well as large quantities of data have contributed to the drastic development of the field. However, the application of deep learning is often hindered by the need for big data and the laborious manual annotation thereof. To experience deep learning using the data compiled by us, we collected 2429 constrained headshot images of 277 volunteers. The collection of face photographs is challenging in terms of protecting personal information; we established an online procedure in which both the informed consent and image data could be obtained. We did not collect personal information, but issued agreement numbers to deal with withdrawal requests. Gender and smile labels were manually and subjectively annotated only from the appearances, and final labels were determined by majority among our team members. Rotated, trimmed, resolution-reduced, decolorized, and matrix-formed data were allowed to be publicly released. Moreover, simplified feature vectors for data sciences were released. We performed gender recognition by building convolutional neural networks based on the Inception V3 model with pre-trained ImageNet data to demonstrate the usefulness of our dataset.

## Introduction

Deep learning is often viewed as having been catapulted into fame by the significant achievements of AlexNet at the ImageNet Large Scale Visual Recognition Challenge in 2012 (Zahangir Alom et al., 2018). AlexNet is a deep convolutional neural network (CNN) consisting of convolutional, max-pooling, fully connected, and softmax layers (Krizhevsky et al., 2012). Long before this well-known breakthrough, a CNN with a back-propagation algorithm, namely LeNet, was proposed and implemented for the recognition of handwritten digits (LeCun et al., 1998). With the advancements in computer hardware performance, CNNs have become popular in computer vision as one of the most successful artificial intelligence methodologies (LeCun et al., 2015).

In addition to the progress made in hardware and software, the contribution of big data to the success of deep learning has been substantial. Deep learning is based on a supervised machine learning algorithm that relies on a large quantity of labeled data. More than one million labeled photographs from ImageNet were used for AlexNet in 2012 (Krizhevsky et al., 2012). ImageNet is a dataset consisting of quality-controlled photos organized into more than 20,000 categories in a hierarchical manner (Russakovsky et al., 2015). It covers up to 15 million images, each of which has been hand-annotated. Without such collection and annotation, deep learning would not have progressed to the current level.

In the early days of this field, face recognition and gender classification tools were dependent on soft biometric traits that could be perceived by humans. SEXNET, which was the first neural network for gender classification, achieved accuracy of over 90% (Golomb et al., 1990) although it learned only 90 facial images at that time. Since then, numerous facial datasets, such as FERET (Phillips et al., 1998), LFW (Huang et al., 2008), Adience (Eidinger et al., 2014), and UTKFace (Zhang et al., 2017), have been developed and released. The majority of these datasets imposed challenging tasks when using images of an unconstrained nature. Whereas headshot data can readily be obtained from the Internet, formally collecting such headshots with agreement from each subject is a formidable task regarding the management of personal information. In contrast to unconstrained images, constrained images may be analyzed to understand the unrecognized soft biometric traits of human faces.

In the current study, we first formed a study group for the data sciences, traversing many departments and backgrounds in our institute, the National Center for Child Health and Development (NCCHD). We endeavored to construct an online collection system of headshots and to release a dataset of annotated headshots for use in deep learning. The dataset includes not only two-dimensional images but also one-dimensional feature vectors that are beneficial for other data sciences. After agreement was obtained from the volunteers, the system requested them to upload image data without personal information. When photographs were captured, the pose and action were constrained. Deviated headshots were manually eliminated, or were rotated, scaled, and set in fixed positions. After reviewing each face, we voted on the gender and smiling labels, which were then determined by majority. By employing leave-one-out cross-validation on all of the released data, our CNN could achieve an accuracy of 98.2% in recognizing the gender of a person.

## Results

### Application to ethics review committee

Our initial members intended to perform deep learning by using our collected data. The idea was to surmise the gender from a headshot image or face photograph. We were interested in determining how accurately common deep learning methods could recognize the gender without individual bearings; that is, by using constrained photographs. When we consulted on this matter with the director of our research institute on July 5, 2018, he encouraged us to proceed and advised us to begin with applying for ethics review. The idea was approved with a number of 2045 on January 10, 2019. However, there were several restrictions. We decided not to collect the volunteer names, gender, and contact information including e-mail addresses. We provided the volunteers with a withdrawal right for 50 days after uploading their photographs. We were allowed to publicly release the data after trimming, size and resolution reduction, decolorizing, and converting them into array forms consisting of only pixel intensity data.

### Preparation for data collection

The initial team constructed a CGI submission system to obtain not only informed consent but also headshots from the volunteers. Our intention and purpose appeared first on the website, following which we clarified what we would and would not collect. Through this process, we also collected the submission timestamps, IP addresses, and original file names from the submitters for the purpose of identifying the volunteers. Volunteers were requested to straighten up, look in the front, take off glasses, hats, scarfs, mufflers, masks, and so forth, and use plain background for the photographs as much as possible. Make-up including wearing earrings was permissible. They were requested not to cover eyes with hair and not to cover their faces. The first inquiry confirmed whether he or she was ready to take his or her 9 photographs or already had 9 photographs captured for this purpose. We did not accept photographs taken for other purposes. The second confirmed whether he or she was the person concerned. The third confirmed whether he or she understood our motive and purpose, and agreed to submit photographs of their own. The fourth confirmed the conditions for the withdrawal of the agreement. Thereafter, the year of birth, with neither the month nor day, was required to confirm that the volunteer was no less than 20 years old. We did not collect data from individuals whose age was less than 20 years.

### Collection of volunteers and their headshot images

We first sent e-mails to all employees working at our institute, the NCCHD, on February 7, 2019. The first volunteer submitted his or her data the following day. Thereafter, we hosted several explanatory seminars and announced the recruiting of not only volunteers who could generously provide their headshot images, but also team members who wished take part in the study of data sciences. By the end of January 2020, our web server had received 282 submissions. We requested 9 face images per volunteer and obtained all images from 279 people, although we failed to obtain 1 and 9 images from one and two volunteers, respectively. All of the successfully uploaded image files were in JPEG format. The smallest image file was 54.2 KiB with 960 × 721 pixels, whereas the largest one was 5.03 MiB with 4160 × 3120 pixels. We used the Exif Orientation information and did not look at other metadata. The 9 uploaded files were investigated using a POSIX command cmp repeatedly to determine whether identical files existed. The redundancy of these files was eliminated by removing either of the identical files. Several submitters uploaded identical files twice or more in the 9-file set. Some complained to us that 9 image files were too many to handle using a smartphone. The calling for volunteers was not restricted to employees or to Japanese. Nevertheless, the majority of the volunteers appeared to be Japanese or East Asian. Although individuals who appeared South Asian or European were included, our collection was biased in terms of skin-type or racial balance (Buolamwini and Gebru, 2018). Instead of collecting personal information, we instantly issued an agreement number to each volunteer to deal with subsequent withdrawal requests. However, we did not receive any such requests or inquiries.

### Quality control of uploaded image files

When the team members flicked through the images, we noticed that one volunteer uploaded his or her data thrice and another uploaded his or her data twice. The second and third redundant submissions were deleted. We first employed OpenCV face recognition to prepare the labeling and training data, and although it functioned acceptably in most cases, we opted to rescue rare failure cases. Therefore, we manually drew a rectangle that fit each facial contour individually using the image manipulation software GIMP. For all of the images, the positions of the rectangles were subsequently saved as a tab-delimited text file to be processed with OpenCV. Several images were eliminated during this process as they did not meet our requirements. These included images with hands and in which the person was not looking in the front. Although we requested the volunteers to remove their glasses, a few volunteers uploaded images with glasses. However, as we felt that the glasses did not affect the classification substantially during the review, we did not eliminate these images. Inclined faces were modified by rotating the entire image to make it upright by using the image manipulation software ImageMagick. The contour positions were obtained from the rotated images. Although very light and dark images existed, the brightness was not modified at all. Finally, 2429 headshot images were obtained from 277 volunteers. For a certain volunteer, only 5 headshots among the 9 uploaded remained, and these were released. Likewise, 6, 7, and 8 uploaded images remained for 4, 11, and 26 volunteers, respectively. For the other volunteers, all 9 images were released, resulting in a total of 2429 images.

### Gender labeling for each volunteer and smile labeling for each image

Because we focused on the appearance rather than biological gender, we did not collect any data regarding gender from the volunteers. Instead, we constructed an in-house voting system to create gender and smile labels. All thumbnails were ordered and displayed using HTML5 to be labeled by the team members. Labeling was carried out by clicking radio buttons and the CGI program subsequently created a voting sheet from each member. All voting sheet sets were finally tallied for the labeling. Therefore, once the subjective voting was completed, the final labeling was objectively completed by computer programs based on majority.

One member was aware that his male friend had provided his images by cross-dressing with make-up. Judging from the appearance in the uploaded images, the member voted for female regarding the male volunteer, without sharing his knowledge with the other members. Because almost all members clicked the female button, the volunteer was labeled as female. This was one of the occurrences during our collecting and labeling processes. However, we did not reveal the image IDs related to these. Among the 277 volunteers, 130 and 147 were labeled as male and female, respectively (Fig. 1).

**Figure 1.**
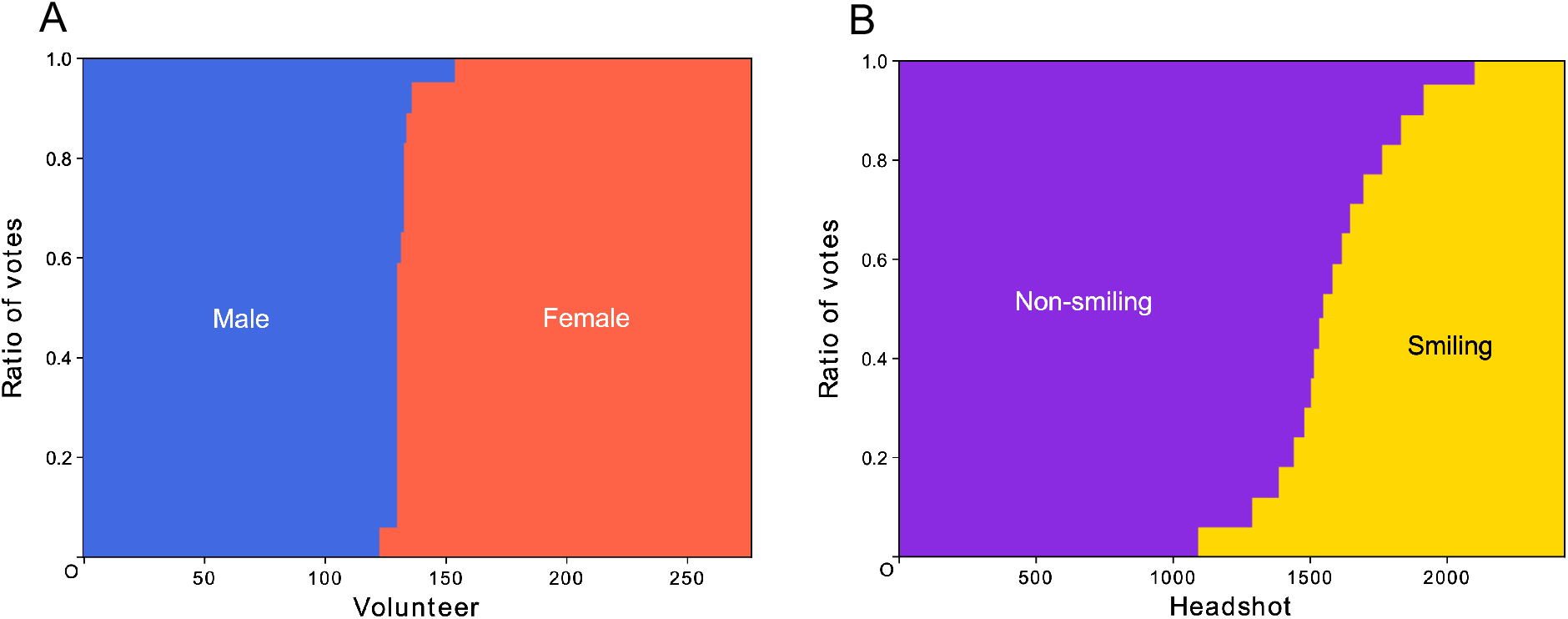
Results of voting for gender and smile labeling. All 277 volunteers and all 2429 headshots were voted on by the team members for the gender and smile labeling, respectively. They were sorted according to the results and renumbered on the horizontal axis. Votes were broken for several volunteers numbered around 130 (A) and for the headshots numbered approximately 1400–1800 (B).

In addition to gender labeling, we endeavored to label faces with smiling. However, determining whether the person was smiling was very difficult. In hundreds of cases, the team members hesitated to click the radio button. However, the voting system that adopted majority mitigated the manual labeling process. Unlike the gender labels, smiling was indicated by the numbers of votes obtained, such as 10 and 7 for non-smiling and smiling, respectively. The total number of voters was limited to an odd number of 17 (Fig. 1).

### Two-dimensional and one-dimensional data

Based on the rectangles that fit each upright facial contour, we cropped the images and resized them to prepare 359 × 359 array data for CNN. The faces were decolorized and placed in the center with enough margins. For example, if the centers are cropped into 299 × 299 pixels and expanded to three channels, they can be readily used as input data for the Inception V3 network.

During the data processing, we observed that the gender could simply be distinguished by the existence of long hair around the level of the mouth or jaw in various cases. For the convenience of applying these to other machine learning methodologies, simplified one-dimensional data were also produced, which consisted of one feature vector per volunteer and 75 features derived from horizontal pixel intensities at the jaw level. Bilateral symmetric and neighboring positions were merged to form the 75 features. The merged intensity values were min-max normalized and arranged from the outside to the center. Hierarchical clustering of the feature vectors was carried out to visualize the values, which suggested the existence of dark pixels around the middle of the array in numerous female samples (Fig. 2). Although principal component analysis could roughly separate the male and female samples, its decomposition ability was far from acceptable (Fig. 2). In addition to these unsupervised learning methods, we carried out linear classification, random forest, and gradient boosting using the feature vectors. The accuracies obtained by the leave-one-out cross-validations were 76.5%, 81.2%, and 80.1%, respectively. The top five that contributed to the classification were the 40th, 46th, 42nd, 44th, and 49th features. These were numbered using a 0-based method.

**Figure 2.**
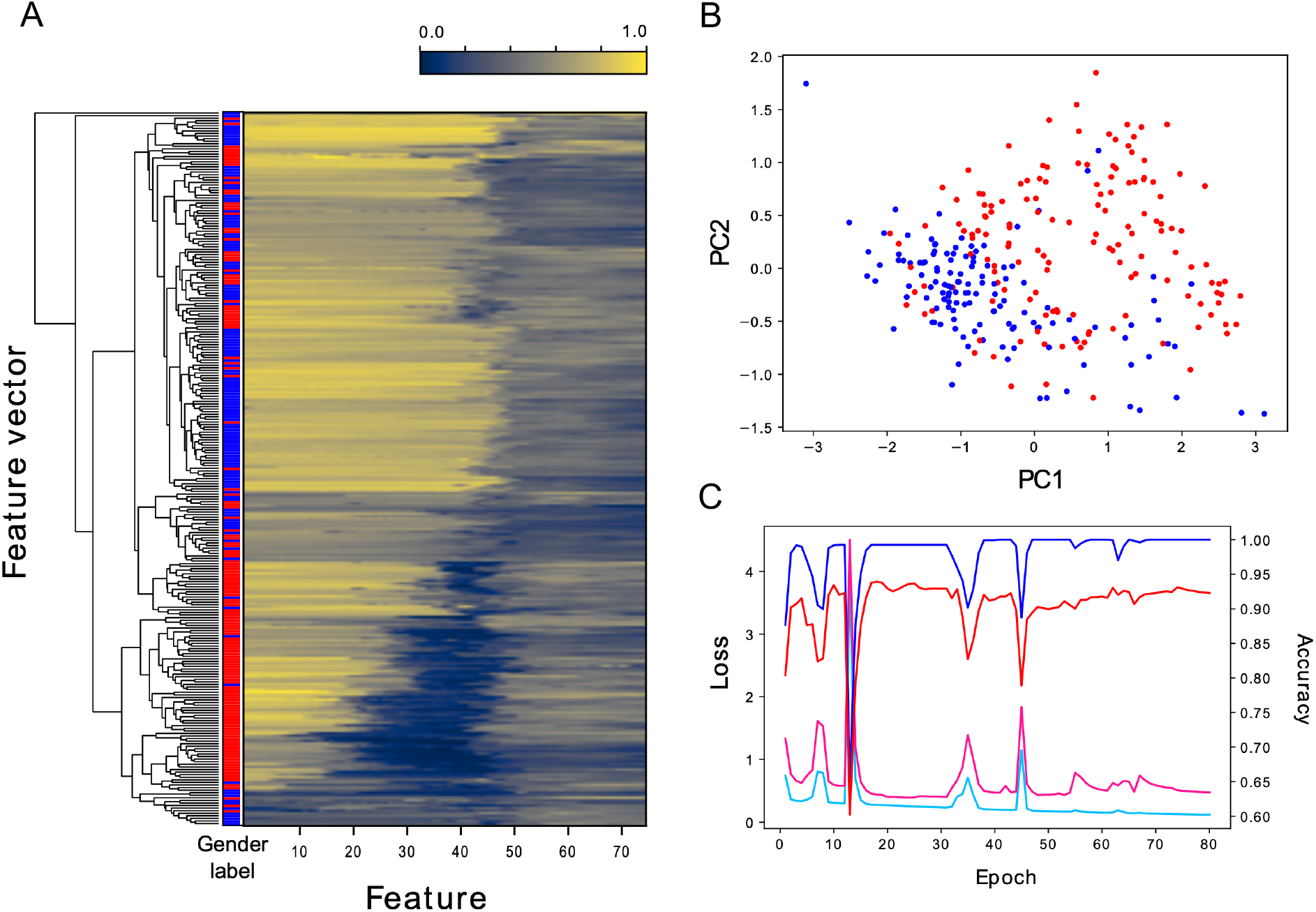
Hierarchical clustering, principal component analysis, and deep learning. Hierarchical clustering was performed for the mandible feature vectors, which consisted of 277 samples and 75 features (A). The darker color corresponds to the darker parts in the headshot image. Although genders were not separated effectively by this unsupervised method, female samples tended to exhibit darker parts around the halfway point. Principal component analysis was performed for the feature vectors (B). The blue and red dots represent 130 male and 147 female samples, respectively. Although they preferentially occupy the bottom left and top right corners, respectively, the two gender groups could not be clearly discerned by this unsupervised method. Learning curves of the CNN based on the Inception V3 is shown (C). Since the leave-one-out cross-validation with 10-times augmentation could achieve high accuracy in a single epoch, learning curves of a 2-fold cross-validation without data augmentation are shown. Odd-numbered and even-numbered volunteer samples were used as training and test data. Exemplary curves were observed during the first 5 epochs in this case. Cyan, pink, blue, and red lines indicate training loss, test loss, training accuracy, and test accuracy, respectively.

Finally, using the two-dimensional data of the 2429 images, we performed gender classification by deep learning (Fig. 2). In the leave-one-out cross-validation, all headshot data up to 9 from one volunteer remained and were used as test data. When we adopted Inception V3 with the ImageNet pre-trained data, the network guessed the right gender for 272 volunteers, with an accuracy level of 98.2%.

## Discussion

Over the course of approximately one year, we recruited more than 200 volunteers online. The campaign was not straightforward. We initially anticipated that young people would be interested and advertised the initiative through several social networking services. However, contrary to our expectations, the number of volunteers barely increased until we held recruiting seminars repeatedly. Calling out to the employees at our institute also contributed to the final quantity of the collected data. The online collection system required each volunteer to validate his or her year of birth to confirm that they were no younger than 20 years old. Although we neither used the data for deep learning nor released the data, the percentage of volunteers that were less than 30 years old was 14.8%. It is likely that older people tended to be more lenient in providing their own photographs.

The campaign was challenging owing to the current trend of protecting personal information. Prior to collecting the data, we consulted with members of the ethics review committee several times to determine what was and was not allowed. Although our proposal using the online system was finally approved, we had to compromise by limiting the types of collected data. It is unfortunate that we could not release colorized or three-channel data. However, the data quantity would have been much lower if we had not decolorized the images. We have demonstrated that using all these volunteer data is sufficient to predict their gender using CNN. Many congenital disorders such as Down syndrome, Noonan syndrome, and Williams syndrome can be diagnosed to some extent from their characteristic facial features by experienced doctors (Gurovich et al. 2019). Since most of them are rare diseases, collecting data would be a demanding task. However, our results encourage us to proceed with such tasks. We are still seeking more sophisticated means of collecting even more constrained facial data with informed consent using other technologies, such as blockchain.

Although requesting 9 photographs appeared to place a heavy burden on volunteers, many images of a single person aided us in creating labels unexpectedly. This was applicable to both the gender and smile labeling. In certain female cases, it was difficult to draw conclusions using a single photograph. Reviewing several facial expressions may play a role in the recognition of a person. As the selection of smiling faces was rather subjective, many images were certainly necessary. Because most photographs were taken in “selfie” modes, natural smiling did not appear frequently in our collections. Constrained headshots were not suitable for smile labeling.

Several studies have concluded that face recognition is more accurate in men than in women (Buolamwini and Gebru, 2018; Albiero et al., 2020). Accordingly, our CNN committed errors in 2 male and 3 female cases. All 3 females had short hair; one of the 2 males had long hair. Compared to unconstrained datasets, our collection was relatively small. However, as a constrained headshot dataset with informed consent and hand-annotations, the scale was unprecedented. Resolution of each image was much higher than those obtained by web scraping. The several images, up to 9, from each individual was also a significant advantage. We could straightforwardly observe the importance of hair length, which was also supported by the one-dimensional mandible vectors. Despite the fact that our collection system was initially built for educational purposes, it appeared to offer the possibility of identifying unrecognized biometric traits in the faces.

## Methods

### Ethics approval and protection of personal information

All methods were performed in accordance with the relevant national guidelines and regulations. The present study was approved by the NCCHD ethics review committee, approval number 2045 on January 10, 2019. As described in the result section, informed consent was obtained online, which was also approved by the committee.

### Hardware and software

We shared NEC Express5800/R120d-2E in addition to individual personal Linux, macOS, or Windows computers. We used Hitachi HA8000/RS210, HPE ProLiant DL360, and Hitachi SR24000/DL1 for the high-intensity calculations. For data collection, data release, and demonstration of our own application, we used hosting services with global IP addresses. We made use of numerous online services, including FC2 Wiki, Slack, Facebook, GitHub, and Google Docs, without any payments. For the most part, we used CentOS 7.6, but we attempted various Linux distributions, from Ubuntu and openSUSE to Windows Subsystem for Linux. We exclusively employed free software or open-source software as far as possible, including Anaconda, TensorFlow, Keras, OpenCV, GIMP, ImageMagick, Jubatus, Scikit-learn, and LightGBM.

### Headshot data and labels

The shape of the released images was a 359 × 359 pixel square, which was reshaped into a 128,881-dimensional vector row by row. To ensure the equality of the volunteers, the vectors were shuffled and concatenated into an array saved as a NumPy file. The presented order could be retrieved from the bundled label data, which were provided as another NumPy file. The file consisted of five columns: volunteer ID, photograph number for a given volunteer, female label, non-smile label, and smile label. The volunteer IDs were serial numbers from 1 to 277. The photograph numbers ranged from 1 to 9, but there were missing numbers for certain volunteers. If the volunteer was considered a female by our decision rule, the column was labeled as 1. Non-smiling and smiling labels were indicated by numbers of votes.

### Mandible feature vectors

In addition to the headshot data, a simplified dataset was provided. The data were derived from horizontal pixel intensities around the level of the mouth or jaw, in the 267th, 268th, and 269th rows from the top, excluding 30 pixels on both sides. These were numbered with a 0-based method. The 6 neighboring pixels were merged, except for the 9 center pixels. Furthermore, bilateral symmetric positions were merged to form the 75 features or explanatory variables. The merged intensity values were min-max normalized, from 0.0 to 1.0, and arranged from the outside to the center around the mouth.

### Unsupervised and supervised machine learning

Hierarchical clustering was carried out using the Euclidean distance and average linkages. Clustering and drawing of the heatmap and dendrogram were achieved with the SciPy library 1.1.0, https://www.scipy.org/scipylib/. Principal component analysis was carried out using the Scikit-learn library 0.21.2, https://scikit-learn.org/. Linear classification was performed using Jubatus 1.1.1, http://jubat.us/, with the AROW algorithm, a regularization weight of 1.0, and 100 epochs. Random forest was applied using the Scikit-learn library with a maximum depth of 32, 16 trees, and a random state of 32. Gradient boosting was carried out using LightGBM 2.3.0, https://github.com/microsoft/LightGBM. Binary log loss classification was selected with 38 boosting rounds. If a predicted value was no less than 0.5, the sample was inferred as female. Inception V3 with the ImageNet pre-trained data yielded the optimal results for the CNN gender classification. The network was implemented in Keras 2.2.4, https://keras.io/, with TensorFlow 1.12.0, https://www.tensorflow.org/, as a backend. We added three fully connected layers and a sigmoid layer for the binary classification. We set the 225^th^ and latter layers to be trainable. The training data were augmented 10 times using Keras ImageDataGenerator with a rotation range of 6.0, width shift range of 0.06, height shift range of 0.06, shear range of 2, zoom range of 0.08, and horizontal flip on. The batch size and number of epochs were 64 and 20, respectively. Our Python scripts are available at https://github.com/glires/Foxglovetree/.

### Availability of data and programs

The image data and labels, namely the Foxglovetree dataset, are publicly available from the following sites, https://aihospital.ncchd.go.jp/foxglovetree/data/ or https://github.com/glires/Foxglovetree. In appreciating the generosity of many volunteers, we allow neither distribution nor release of any image formats, such as JPEG files, derived from our data. The scripts used to process the data and to perform machine learning are also available at https://github.com/glires/Foxglovetree or our portal site, https://aihospital.ncchd.go.jp/foxglovetree/.

## Acknowledgements

Our small group activity was originally encouraged by the director of the research institute, Dr. Yoichi Matsubara, and subsequently by the NCCHD AI hospital project, particularly by Dr. Yasuo Kiryu. Ms. Kyoko Hagino provided us with constructive suggestions regarding the protection of personal information. Although we did not receive any financial support for the activity itself, the publication cost was supported by the AI hospital project, officially, “Innovative AI Hospital System”, Cross-ministerial Strategic Innovation Promotion Program (SIP), Council for Science, Technology and Innovation (CSTI). The National Institute of Biomedical Innovation, Health and Nutrition (NIBIOHN) is its funding agency. It was also supported by Takeda Science Foundation. We did not purchase any hardware for this study. However, several departments and individual employees generously provided used computers. Our main machine, NEC Express5800/R120d-2E, was provided by BioBank, which is directed by Dr. Kenichiro Hata. We also used the Hitachi HA8000/RS210 and HPE ProLiant DL360 clusters at the Center for Regenerative Medicine, which is directed by Dr. Akihiro Umezawa, Hitachi SR24000/DL1 P100 purchased by Dr. Yoichi Matsubara, and the same model with V100 purchased by the AI hospital project. Finally, we sincerely appreciate all of the volunteers for providing their headshot image data.

## Author contributions

KO conceived and designed the study. YI applied for ethics review. SA, MH, HU, AU, MI, YI, MT, TJ, MS, YO, FH, HO, SN, KK, MK, KM, KT, and KO collected, curated, annotated, and analyzed the data and discussed the results. KO compiled the release data. TM, KT, and KO drafted the manuscript.

